# Theta-phase connectivity between medial prefrontal and posterior areas underlies novel instructions implementation

**DOI:** 10.1101/2022.02.23.481594

**Authors:** Silvia Formica, Carlos González-García, Mehdi Senoussi, Daniele Marinazzo, Marcel Brass

## Abstract

Implementing novel instructions is a complex and uniquely human cognitive ability, that requires the rapid and flexible conversion of symbolic content into a format that enables the execution of the instructed behavior. Preparing to implement novel instructions, as opposed to their mere maintenance, involves the activation of the instructed motor plans, and the binding of the action information to the specific context in which this should be executed. Recent evidence and prominent computational models suggest that this efficient configuration of the system might involve a central role of frontal theta oscillations in establishing top-down long-range synchronization between distant and task-relevant brain areas. In the present EEG study (human subjects, 30 females, 4 males), we demonstrate that proactively preparing for the implementation of novels instructions, as opposed to their maintenance, involves a strengthened degree of connectivity in the theta frequency range between medial prefrontal and motor/visual areas. Moreover, we replicated previous results showing oscillatory features associated specifically with implementation demands, and extended on them demonstrating the role of theta oscillations in mediating the effect of task demands on behavioral performance. Taken together, these findings support our hypothesis that the modulation of connectivity patterns between frontal and task-relevant posterior brain areas is a core factor in the emergence of a behavior-guiding format from novel instructions.

**Significance statement:** Everyday life requires the use and manipulation of currently available information to guide behavior and reach specific goals. In the present study we investigate how the same instructed content elicits different neural activity depending on the task being performed. We show that medial prefrontal theta oscillations are larger when novel instructions have to be implemented, rather than simply maintained. Crucially, connectivity between medial prefrontal cortex and posterior areas is strengthened during instructions implementation, suggesting that theta oscillations play a role in setting up a dynamic and flexible network of task-relevant regions optimized for the execution of the instructed behavior.

## Introduction

The ability to rapidly adapt to changing external contingencies is a crucial hallmark of human cognition. This flexibility finds one of its most advanced and astonishing forms of expression in instructions following. Humans can implement novel behaviors based on symbolic instructions, in the absence of prior practice (Cole et al., 2013). For instance, the driver reaching unexpected roadworks might be presented with different options on how to proceed. They will have to encode and understand their content and execute one of them at the appropriate moment (e.g., “To reach the train station, take the second exit at the roundabout”). This apparently simple operation involves many complex cognitive processes, ultimately resulting in the execution of the correct behavior.

It is established that maintaining task information is not sufficient to successfully perform the instructed behavior (Milner, 1963; Duncan et al., 1996; Bhandari and Duncan, 2014). Rather, the abstract content of the instruction needs to be reformatted into an action-oriented code capable of driving behavior (Brass et al., 2017). At the hemodynamic level, several fMRI studies revealed a set of frontoparietal regions recruited to support novel stimulus-response mappings (SRs) implementation and showing representational patterns specific to execution task-demands (Ruge and Wolfensteller, 2010; Demanet et al., 2016; González-García et al., 2017, 2021; Muhle-Karbe et al., 2017; Bourguignon et al., 2018; Palenciano et al., 2019a, 2019b; Ruge et al., 2019). Although evidence is accumulating concerning the neural substrate deployed for these reformatting purposes, the mechanisms binding stimulus and response information in an action-oriented format, and its exact nature, remain elusive.

In a recent EEG study, we investigated the differences in oscillatory activity associated with maintenance and implementation of novel instructions (Formica et al., 2021). The two tasks showed analogous attentional prioritization mechanisms, reflected in suppression of alpha power contralateral to the attended location (Jensen and Mazaheri, 2010; Gould et al., 2011; Bonnefond and Jensen, 2012; Myers et al., 2015; Mok et al., 2016). On the contrary, proactively transforming the content of the mapping into an action-oriented representation (i.e. the implementation task) elicited task-specific cognitive processes. Namely, preparing a defined motor plan was associated with stronger engagement of motor areas, indexed by beta suppression over motor and premotor cortices (Cheyne, 2013; Schneider et al., 2017). Furthermore, increased theta power over midfrontal scalp electrodes during implementation was interpreted as a marker of working memory (WM) manipulation (Onton et al., 2005; Itthipuripat et al., 2013) and top-down control over alternative task-sets (Miller and Cohen, 2001; Sauseng et al., 2010).

Notably, midfrontal theta oscillations received increased attention in recent years due to their occurrence across a variety of tasks characterized by need for cognitive control (Cohen and Donner, 2013; Cavanagh and Frank, 2014). Theories and evidence emerged regarding slow oscillations as prime mechanism for establishing top-down neural communication and for efficiently setting up functional networks optimized to meet task demands (Sauseng and Klimesch, 2008; McLelland and VanRullen, 2016; Voloh and Womelsdorf, 2016; Bonnefond et al., 2017; Riddle et al., 2020a, 2020b). In line with this framework, recent computational models have attributed to theta oscillations a crucial role in flexibly binding posterior task-relevant areas (Verguts, 2017; Senoussi et al., 2020b; Verbeke et al., 2020). Burst of theta-locked activity originating in the medial prefrontal cortex (mPFC) are directed towards the appropriate processing units (i.e., areas encoding task-relevant information), inducing the synchronization of their firing patterns and thus enabling efficient inter-areal communication (Fries, 2005, 2015). However, the specific role of theta oscillations in instruction implementation remains untested.

Here, we set out to investigate the hypothesis that medial prefrontal theta oscillations not only differ across task-demands (i.e., maintaining vs implementing novel SRs), but also exert different influence on posterior task-relevant areas through long-range connectivity, reflecting the need to orchestrate the synchronization between stimulus and response information to form action-oriented representations.

## Methods

### Participants

Thirty-five participants were recruited from the experimental pool of Ghent University and gave their informed consent at the beginning of the experiment, in accordance with the ethical protocols of Ghent University. Eligibility criteria included age between 18 and 35 years and no history of psychological or neurological conditions. Sample size was not computed a-priori, but chosen to replicate the results of a previous study with a similar paradigm (Formica et al., 2021) and in line with the literature on similar constructs (Schneider et al., 2017; de Vries et al., 2018; van Ede et al., 2019). The experiment consisted of two separate sessions, one to three days apart from each other. In the initial behavioral screening session (∼30 minutes) participants practiced the two tasks, starting always with the Implementation task followed by the Memorization task. Initially, they could familiarize with the task requirements by performing without time pressure (i.e., no response deadline). After responding correctly to 5 consecutive trials, the normal 2000 ms response deadline was introduced. Participants then completed 4 mini blocks of 15 trials each, receiving feedback after each response and at the end of the block as percentage of accurate responses. If overall performance in the 60 trials reached an accuracy threshold of 70% in both tasks, the participant was invited to take part in the EEG session, otherwise they would be compensated for the behavioral screening (5€). Participants could repeat the practice blocks once if they did not reach the threshold at their first attempt. Two participants repeated the practice for the Memorization task, and one participant for the Implementation task. Eventually, all participants reached the required threshold in both tasks. One participant dropped out after completing the behavioral screening, leaving a sample of thirty-four participants in the main EEG session (M_Age_ = 21.73, SD_Age_ = 3.29, 30 females, 4 males). All participants had normal or corrected-to-normal vision and twenty-nine reported to be right-handed. Data from three participants were discarded because of low task performance (their individual accuracy exceeded by 2.5 standard deviation the mean group accuracy and/or their accuracy was below 60% in response to catch trials in at least one of the two tasks). One additional participant was discarded because of noisy EEG recordings. Therefore, our final sample consisted of thirty participants.

### Materials

The set of stimuli consists of three macro-categories: animate (non-human animals), inanimate 1 (vehicles and musical instruments), inanimate 2 (household tools and clothes), each with approximately ∼700 items (Griffin et al., 2007; Konkle et al., 2010; Brady et al., 2013; Brodeur et al., 2014; González-García et al., 2020). All pictures were centered in a 200x200 pixels canvas, were converted to grayscale, and had their background removed.

### Procedure

Stimuli presentation and response collection were performed using the Psychopy toolbox (Peirce, 2007). In the EEG session, participants performed both tasks (i.e., Implementation task and Memorization task), in a blocked design, with task order counter-balanced across participants. Trial structure was identical in the two tasks, except for the probe screen (Figure 1). Each trial started with a red fixation cross presented in the middle of the screen for 2000 ms (±100 ms), indicating the inter-trial interval. Next, four SRs were presented simultaneously for 5000 ms, one for each quadrant of the screen. Each mapping consisted of one image and one digit from 1 to 4, corresponding to the four response options (1: right middle finger, key “e”; 2: right index finger, key “r”; 3: left index finger, key “i”; 4: right middle finger, key “o”). Importantly, responses could be organized according to two response schemes. Namely, mappings involving a response with the left(right) hand could be presented on the left(right) side of the encoding screen; or they could be presented on the right(left) side of the encoding screen, the latter case resulting in an incongruency between location on the screen and response hand. This was done to orthogonalize the presentation side on the encoding screen and the required response hand. To simplify the encoding of the four mappings, index fingers responses were always associated with the two upper images, and middle fingers responses with the lower images. Each encoding screen contained two images from two different categories, grouped on the left and right side of the screen. Each image was presented only once throughout the whole experiment to ensure the novelty of all mappings. After the presentation of the mappings, a fixation interval of 750 ms was presented, considered sufficiently long to prevent iconic memory traces (Souza and Oberauer, 2016), followed by a retro-cue presented centrally for 250 ms. The retro-cue consisted of four colored corners, pointing to the four quadrants previously occupied by the mappings. Each side of the retro-cue was colored in either blue (RGB = [0, 155, 255]) or orange (RGB = [255, 100, 0]). These colors were chosen to equate luminance and saliency, thereby preventing low-level confounds imputable to exogenous attention. Participants were instructed at the beginning of the experiment that one color indicated the two mappings relevant for the task, and the other color pointed to the location of mappings that could be discarded. Color assignment was counterbalanced across participants and remained the same for the whole duration of the experiment. The information provided by this retro-cue allowed participants to select two out of the four initially encoded mappings. The selected mappings always contained images belonging to the same category and involved responses with the index and middle fingers of the same hand. The retro-cue was followed by a cue-target interval (CTI) with a jittered duration of 1750 ms (±100 ms). Next, the probe appeared and remained on screen for a maximum duration of 2000 ms or until button press. If no response was provided within the response deadline, a message appeared encouraging the participant to take a short break if needed, and to press the spacebar to resume the task. In the Implementation task, the target consisted of a choice-reaction task, with one of the selected images presented centrally. Participants were asked to press the key with the finger associated to the corresponding mapping. The Memorization task was a delayed match-to-sample task, in which participants were presented centrally with one of the selected images and a response digit. This target mapping had to be compared with the encoded ones, to verify if the image-response association was correct. Participants provided their response by pressing with the finger corresponding to the tick sign (✓) or x sign (☒) displayed at the bottom of the probe screen, to indicate matching or mismatching mappings, respectively. Notably, the assignment of and ☒ locations with respect to the four fingers was randomized on each trial. This task design ensured that the two tasks differed only to the extent a specific response could be proactively prepared during the CTI. In the Implementation task, participants were encouraged to prepare the SRs for execution as soon as the retro-cue indicated the relevant two. On the contrary, the Memorization task only allowed for a declarative maintenance of the two selected mappings, because no action-oriented representation could be built and thus no action plan prepared.

**Figure 1:**
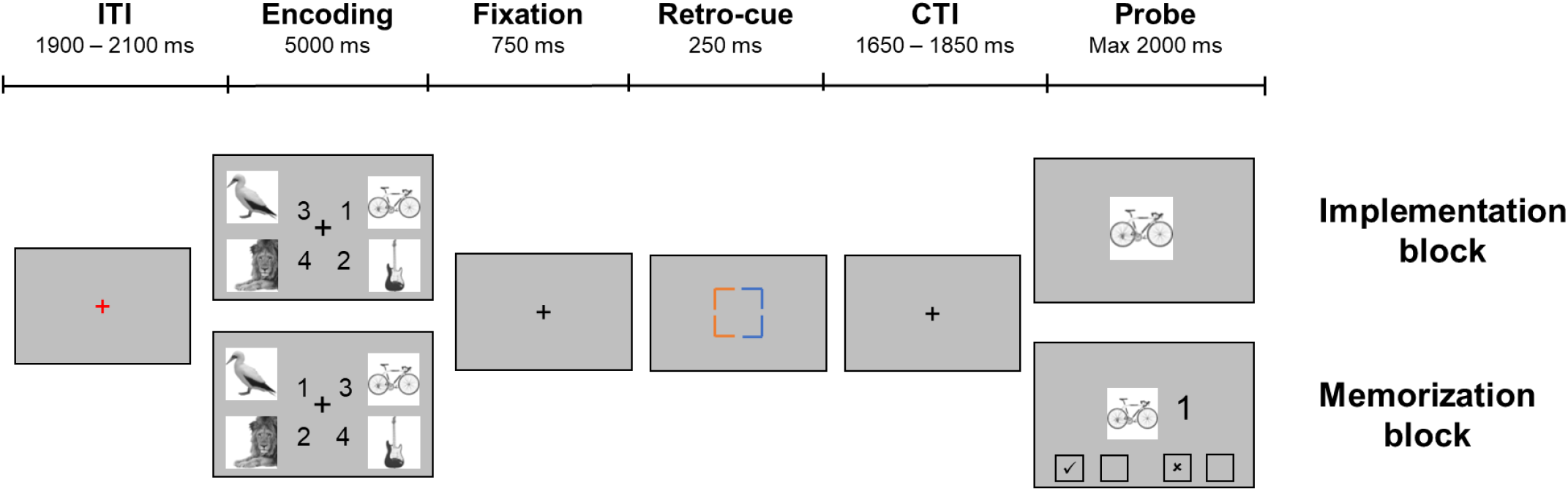
Task paradigm

In both task, 10% of trials featured a completely new image as target, instead of one of the encoded (i.e., catch trials). Participants were instructed to press the spacebar in such events. This was done to ensure all four mappings were initially encoded, discouraging strategies such as focusing only on a subset of mappings. Since we were interested in the brain activity during the CTI, catch trials were included in our analyses.

During the EEG session, 240 trials for each task were completed, divided in 6 blocks. Trials were equally distributed between the 4 conditions resulting from the characteristics of the selected mapping, namely the side they appeared on in the encoding screen (Cued Side, left or right) and the response hand they involved (Response Side, left or right).

### Experimental design and behavioral analysis

The study consisted of three within-subject factors orthogonally manipulated. Namely, Task (Implementation vs Memorization), Cued Side (Left vs Right) referring to the location on the screen of the selected mappings, and Response Side (Left vs Right), indicating the response hand involved in the selected mappings. Based on previous studies comparing implementation and maintenance of novel SRs, we expected to find shorter reaction times (RTs) and lower error rates for the former with respect to the latter (Demanet et al., 2016; Formica et al., 2020, 2021; González-García et al., 2020). Therefore, we used JASP (Jasp Team, 2019) to compare RTs and error rates across Tasks, separately for regular and catch trials. RTs were analyzed by means of paired-samples t-tests. Error rates distributions for both regular and catch trials violated the Normality assumption (Shapiro-Wilk test p < 0.05), and thus the results of Wilcoxon signed-rank tests are reported.

We had no strong hypotheses with respect to the effects of the other experimental manipulations on behavioral performance. In an exploratory fashion, we performed 2 (Task) x 2 (Cued Side) x 2 (Response Side) repeated-measures ANOVAs on RTs and error rates of regular trials. These results are reported in the Supplementary Materials.

### EEG Recordings and pre-processing

Electrophysiological data were collected using a BioSemi ActiveTwo system (BioSemi, Amsterdam, Netherlands) with 64 Ag-AgCl electrodes arranged in the standard international 10-20 electrode mapping (Klem et al., 1999), with a CMS-DRL electrode pair. Two external electrodes were placed on the left and right mastoid for online referencing. Four additional external electrodes (two to the outer canthi of both eyes, one above and one below the left eye) were used to monitor horizontal and vertical eye movements. Data were recorded at a sampling rate of 1024 Hz.

All pre-processing and analyses steps were carried out with the Python MNE toolbox v. 0.22.0 (Gramfort, 2013). First, a 1 – 40 Hz band-pass FIR filter was applied to the continuous data (Hamming window with 0.0194 passband ripple and 53 dB stopband attenuation, with lower and upper transition bandwidth of 1 Hz and 10 Hz, respectively). Next, data were epoched time-locked to the onset of the retro-cue (from -1000 ms to 2500 ms), demeaned to the average of the whole time window to improve independent component analysis (ICA) (Groppe et al., 2009), and downsampled to 512 Hz. Trials containing excessive muscle activity or other clear artifacts were rejected based on visual inspection, and electrodes showing noise for a large portion of epochs were removed and interpolated using the spherical spline method (Perrin et al., 1989). Data were then re-referenced to the average of all channels. Finally, eye movements were removed by means of ICA: an average of 2.23 (SD = 0.716) ICA components were removed for each participant.

Only trials with a correct response were retained for the analyses. This resulted in an average of 218.23 trials for Implementation (SD = 18.89) and 212.70 for Memorization (SD = 12.02). For each individual condition (e.g., SRs presented on the left side of the screen and requiring a left-hand response), an average of 53.86 trials (SD = 4.55) were retained (range 36 – 60 across participants).

### Source reconstruction

To project brain activity from the sensors to the source space, we employed the default anatomical template included in the MNE-Python toolbox (‘fsaverage’ from FreeSurfer) and decimated the dipole grid on the white matter surface to 4098 sources per hemisphere (6^th^ grade subdivisions of an octahedron). A realistic Boundary Element Model (BEM) was created assuming specific conductivity for each of the three shells (skin, outer and inner skull). Noise in the recordings was estimated by computing a noise covariance matrix from the baseline period (-500 to -200 ms) and the inverse problem was then solved with the dynamic statistical parametric mapping (dSPM) method (Dale et al., 2000).

### Regions of Interest (ROIs) definition

To replicate our previous findings and to estimate inter-areal connectivity, we selected an a-priori set of ROIs. From the Desikan-Killiany atlas (Desikan et al., 2006) we obtained left and right lateral occipital parcels (LatOcc ROIs), and left and right caudal anterior cingulate parcels (mPFC ROIs), an area that showed to be consistently activated across a variety of processes for cognitive and adaptive control (Cavanagh and Shackman, 2015; De La Vega et al., 2016). Additionally, we created Hand ROIs by grouping sources within a radius of 30 mm on the inflated surface from the MNI coordinates for the left and right hands ([±44, -17, 49]) (Zhao et al., 2019). The resulting set of ROIs is depicted in Fig 2.

**Figure 2:**
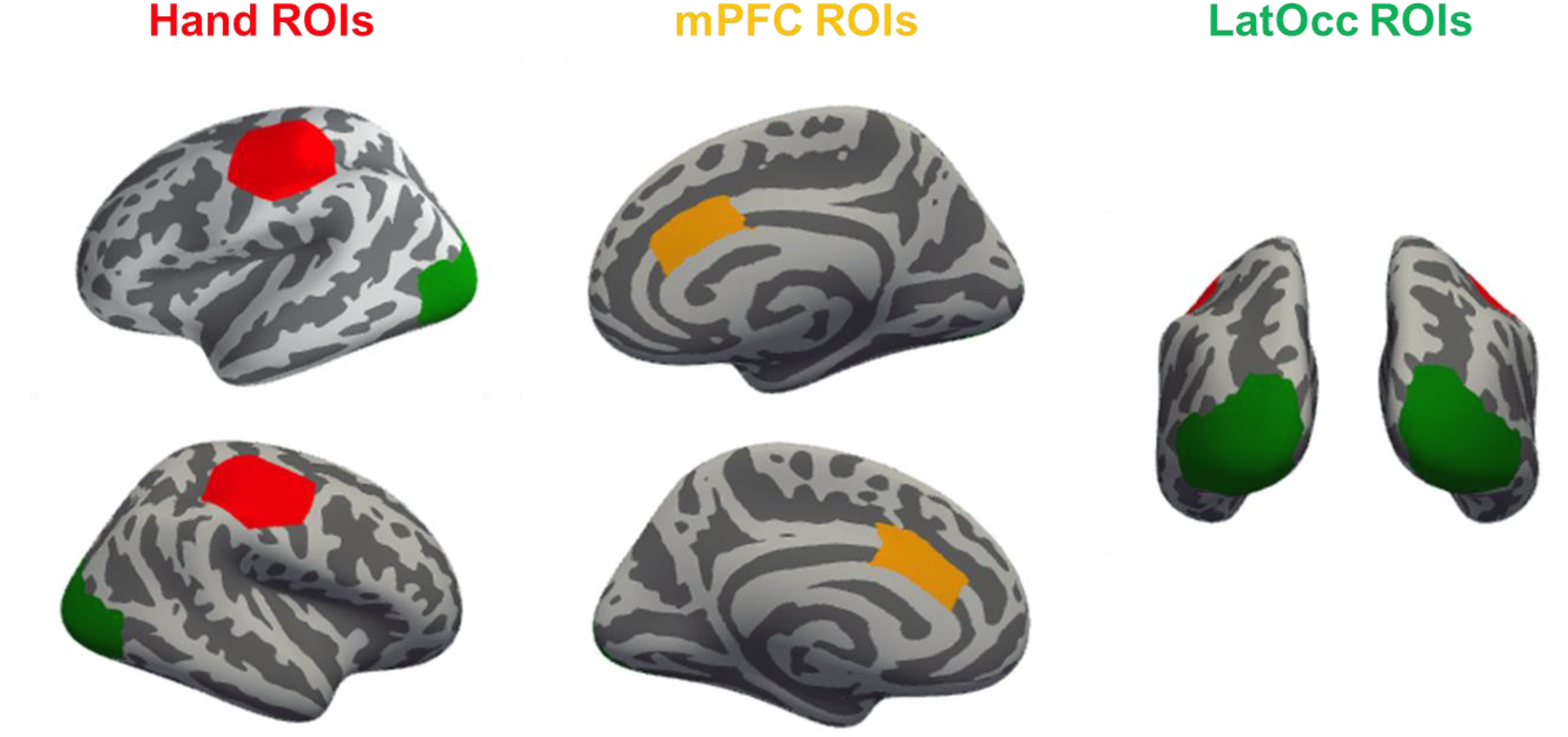
ROIs Locations. mPFC and LatOcc ROIs were obtained from the Desikan-Killiany atlas (caudal anterior cingulate and lateral occipital parcels, respectively). Hand ROIs were drawn with a 30 mm radius around the MNI motor areas hand coordinates ([±44, -17, 49]).

### Statistical Approach

To evaluate the statistical significance of the differences between time courses in our contrasts of interest, we adopted a cluster-based permutation approach (Maris and Oostenveld, 2007). Importantly, this method is effective in evaluating the reliability of neural data patterns over contiguous timepoints, while successfully controlling for the multiple comparisons problem. First, the time courses are compared with an F-test (or t-test, depending on the specific contrast) between all timepoints of the observed conditions, with an α level of 0.05. Next, neighboring time points with above threshold same sign significance are grouped together, and the resulting segment of data is considered a cluster with a size corresponding to the sum of the F-values (or t-values) of all time points belonging to it. The statistical significance of these observed clusters is computed by comparing them to a distribution of clusters obtained under the null hypothesis. Specifically, surrogate data are generated for 10000 permutations by randomly flipping the sign of the difference between conditions. A distribution of cluster sizes under the null hypothesis of no differences between conditions is obtained by taking the size of the largest cluster (computed as for the observed data) for each of the permutations. The P-value for each observed cluster corresponds to the proportion of permutations in which a cluster larger than the observed one was found. Again, we used an α level of 0.05, therefore retaining as significant only clusters larger in size than at least 95% of clusters in the surrogate data. It is important to point out that due to the nature of the second-level inference between observed and surrogate distributions, the exact boundaries of each cluster (i.e., first and last time points) have to be interpreted cautiously and do not reflect the significance of each individual timepoint (Sassenhagen and Draschkow, 2019).

For each observed cluster, an average effect size is computed as Cohen’s d. To obtain it, the difference between the two conditions in the mean values over the time window of significance of the cluster is derived, and then divided by its standard deviation.

All statistical analyses are performed in the time window 0-1800 ms from retro-cue onset. The CTI was jittered, with a duration between retro-cue onset and probe onset ranging between 1900 and 2100 ms. Therefore, the time window for analyses was cut 100 ms before the earliest probe onset, to reduce the influence of the smearing in time resulting from the time-frequency analysis linked to the processing of the probe.

### Spectral Analysis

For each condition, the induced power was computed in the source space for three frequency ranges (theta: 3 – 7 Hz, alpha: 8 – 14 Hz, beta: 15 – 30 Hz). Time-frequency decomposition was performed by means of complex Morlet wavelets, in steps of 1 Hz and with the number of cycles adapted to the frequency range (3 cycles for theta, 4 cycles for alpha, and 5 cycles for beta), to achieve a good trade-off between temporal and frequency precision (Cohen, 2014). The resulting decomposed data was then averaged within each frequency range and further downsampled to 128 Hz.

### Attentional contralateral alpha suppression

When orienting attention to the external or internal space, a suppression of alpha power has been traditionally observed over posterior areas contralateral to the attended hemispace (Jensen and Mazaheri, 2010; Gould et al., 2011; Bonnefond and Jensen, 2012; Myers et al., 2015; Mok et al., 2016). It has previously been shown that Implementation and Memorization do not differ with respect to orienting attention to the relevant encoded mappings (Formica et al., 2021). Here we aim at replicating these findings at the source level. For each condition, time courses of alpha (8 – 14 Hz) power for the left and right LatOcc ROIs were obtained by taking the first right-singular vector of the single value decomposition of all sources within the ROI, scaled to their average power and adjusted in sign for the orientation of each single dipole (‘pca_flip’ method). In other words, this procedure results in a time course explaining the largest possible amount of variance in the whole ROI. The time courses were then collapsed across conditions to obtain an ipsilateral and a contralateral time course with respect to the Cued Side, separately for each of the two tasks. According to our hypotheses and previous findings, we expected to observe significant alpha suppression in the contralateral hemisphere, and no differences between Tasks. To test for this hypothesis, we performed a rmANOVA with factors Tasks (Implementation vs Memorization) and Laterality (ipsilateral vs contralateral with respect to the selected hemispace), adopting a cluster-based permutation approach as described in the Statistical Analysis section.

### Motor contralateral beta suppression

Similarly, beta power is suppressed over the motor cortex contralateral to the limb being prepared for movement (Cheyne, 2013). Time courses of beta (15 – 30 Hz) power activity were extracted from the Hand ROIs ipsilateral and contralateral to the response hand required by the selected mappings, with a procedure analogous to the one used for alpha. Here, we predicted a larger contralateral suppression in Implementation compared to Memorization, due to the proactive preparation of the specific motor plan instructed by the mapping. Therefore, we performed a rmANOVA with factors Tasks (Implementation vs Memorization) and Response Side (contralateral vs ipsilateral to the instructed response hand).

### Task-specific theta increase

Theta power over mid-frontal cortices has been observed to be larger when the SRs are prepared for execution (i.e., Implementation) compared to their declarative maintenance (i.e., Memorization) (Formica et al., 2021). According to prominent computational models, this phenomenon can be interpreted as reflecting the activity of the mPFC in coordinating the neural communication between distant task-relevant areas (Voloh and Womelsdorf, 2016; Verguts, 2017). Therefore, we extracted a time-course of theta (3 – 7 Hz) power activity from the bilateral mPFC ROIs for each of the two Tasks, and we compared, predicting larger values for Implementation.

### Mediating effect of theta power on RTs

We further hypothesized mPFC theta oscillations to play a role in determining behavioral performance. Namely, we expected theta power to mediate the effect of Task on RTs. Trial-by-trial theta power was defined as the average theta (3 – 7 Hz) power in a time window of length 633 ms centered on the cluster of significant difference between Tasks (Task-specific theta increase analysis section). This resulted in an interval spanning from 355 to 985 post retro-cue onset. The choice of this duration is motivated by the need to encompass at least 2 oscillatory cycles at the lowest frequency of interest (3 Hz). Given that single-trial power estimation led to some outliers due to noisy trials, for each participant we removed trials in which the computed mean theta power exceeded 3 standard deviations from the individual mean. Analogously, RTs were trimmed, separately for each Task and participant, removing trials with latencies exceeding 3 standard deviations from the mean^1^. This trimming approach resulted in an average of 6.47 (± 2.11, 1.68%) trials being removed due to outlier values of theta power, 3.1 (± 1.3, 1.58%) and 1.47 (± 1.63, 0.75%) trials being removed with respect to reaction times in the implementation and memorization task, respectively.

To empirically test the prediction that theta power mediates the effect of Task on RTs, we started by verifying the necessary criteria put forward by Baron and Kenny (1986). Namely, it had to be confirmed that Tasks influences both RTs and theta power, and that theta power predicts RTs. We fitted linear mixed effects models (LMMs) using the lme4 package in R (Bates et al., 2014), adopting a backward selection approach to define the random effect structure of the model (Matuschek et al., 2017). The reported P-values were calculated using Satterthwaite approximations (Luke, 2017). Finally, to formally and directly test the mediation effect, a causal mediation analysis was performed with the mediation package in R (Tingley et al., 2014).

### Functional connectivity: Phase Locking Value

According to the computational model proposed by Verguts (2017) and updated by his colleagues (Verbeke and Verguts, 2019; Senoussi et al., 2020b; Verbeke et al., 2020), theta oscillations from the mPFC control unit serve the purpose of synchronizing posterior task-relevant processing units. Therefore, we were primarily interested in investigating whether task-demands (i.e., preparing for implementation vs maintenance) affect connectivity between frontal and posterior areas.

To estimate phase synchronization between pairs of ROIs, we first band-pass filtered our epoched data in the theta frequency range (3 – 7 Hz; Hamming window with 0.0194 passband ripple and 53 dB stopband attenuation, with lower and upper transition bandwidth of 2 Hz). After projection to the source space, data was Hilbert transformed to obtain the analytic signal from which to extract the phase information. Then, we computed the phase-locking value (PLV), a measure reflecting the instantaneous phase difference between two signals, with the assumption that two brain areas that are configured to communicate efficiently should have a constant phase difference (Lachaux et al., 1999; Mormann et al., 2000). Instead of extracting a representative time course for each ROI, resulting in loss of information, we adopted a recently proposed multivariate approach, that allows to efficiently estimate the PLV across all pairwise sources of the two ROIs (Bruña et al., 2018; Bruña and Pereda, 2021). Therefore, the connectivity estimate for each pair of ROIs was obtained by taking the root mean square value of the M x N matrix containing the pairwise PLV between all sources, where M is the number of sources in ROI 1 and N the number of sources in ROI 2.

PLV between pairs of ROIs was computed for each trial in the time window from 355 to 985 ms, resulting in a value of synchronization for each trial, participant, and ROIs pair of interest. It is worth pointing out that PLV is sensitive to volume conduction and source leakage, thus prone to identify spurious connectivity between two neighboring sensors or brain regions reflecting activity from the same source. This concern is moderated in the present study by the selection of ROIs relatively distant from each other, and, most importantly, by comparing connectivity of pairs of ROIs between conditions (see below). Although the magnitude of PLV might theoretically be inflated by source leakage, our hypotheses are focused on how PLV between pairs of ROIs is influenced by different task-demands, and therefore are unaffected by this potential issue.

### Functional connectivity between mPFC and motor and visual areas

We expected the synchronization between frontal and motor areas to be stronger when preparing to implement the SRs, due to the activation and binding of the instructed motor plan. Additionally, we had a further corollary hypothesis, namely that differences between tasks were expected to be driven predominantly by an increase of synchronization between mPFC and the Hand ROI contralateral to the currently relevant response hand in Implementation, but not so in Memorization. In other words, we expected an interaction between the factors Task and Response Side (collapsed for contralateral vs ipsilateral), with a specific directionality. To this aim, we fitted a LMM estimating the trial-by-trial PLV value using Task and Laterality (contralateral vs ipsilateral) as predictors. The model fitting procedure was analogous to the one described above and was conducted using only correct regular trials. The selected model structure included Task, Laterality and their interaction as fixed effects, a random intercept for each participant and a random slope for Task (in lmer notation: *PLV(mPFC-Hand) ∼ Task * Laterality + (1 + Task | Subject)*).

Similar hypotheses and analyses were proposed with respect to connectivity between mPFC and visual areas. The trial-by-trial PLV between mPFC and LatOcc ROIs was fitted with a LMM, predicting an interaction of the factors Task and Laterality (defined as ROI contralateral or ipsilateral to the Cued Side). Model structure was analogous to the one reported for the connectivity between mPFC and Hand ROIs (*PLV(mPFC-LatOcc) ∼ Task * Laterality + (1 + Task | Subject)*).

## Results

### Behavioral results

Based on previous studies reporting performance on implementation and maintenance of novel SRs (Demanet et al., 2016; Formica et al., 2020, 2021; González-García et al., 2021),we expected faster and more accurate responses in the Implementation compared to Memorization task. Indeed, a paired-samples t-test on RTs of regular trials yielded a large effect of Task (*t* _29,1_ = 29.101, *p* < 0.001, Cohen’s d = 5.313), with average RTs in the Implementation task (Mean = 708 ms, ± 129) being shorter than in the Memorization task (Mean = 1217 ms, ± 123). Similarly, a Wilcoxon signed-rank test on error rates showed significantly less errors (*W* _29,1_ = 65.5, *p* = 0.003, Effect size = 0.653) in Implementation (Mean = 0.060 ± 0.047) compared to Memorization (Mean = 0.089 ± 0.029).

RTs and accuracy were compared across Tasks also with respect to catch trials. A paired samples t-test showed support (*t* _29,1_ = 6.508, *p* < 0.001, Cohen’s d = 1.188) for faster RTs in response to catch trials in Implementation (Mean = 817 ms ± 119) compared to Memorization (Mean = 943 ms ± 105). On the contrary, no reliable difference (*W* _29,1_ = 95.5, *p* = 1, Effect size = 0.005) in their accuracy (Implementation: Mean = 0.035 ± 0.070; Memorization: Mean = 0.036 ± 0.061) (Figure 3).

**Figure 3:**
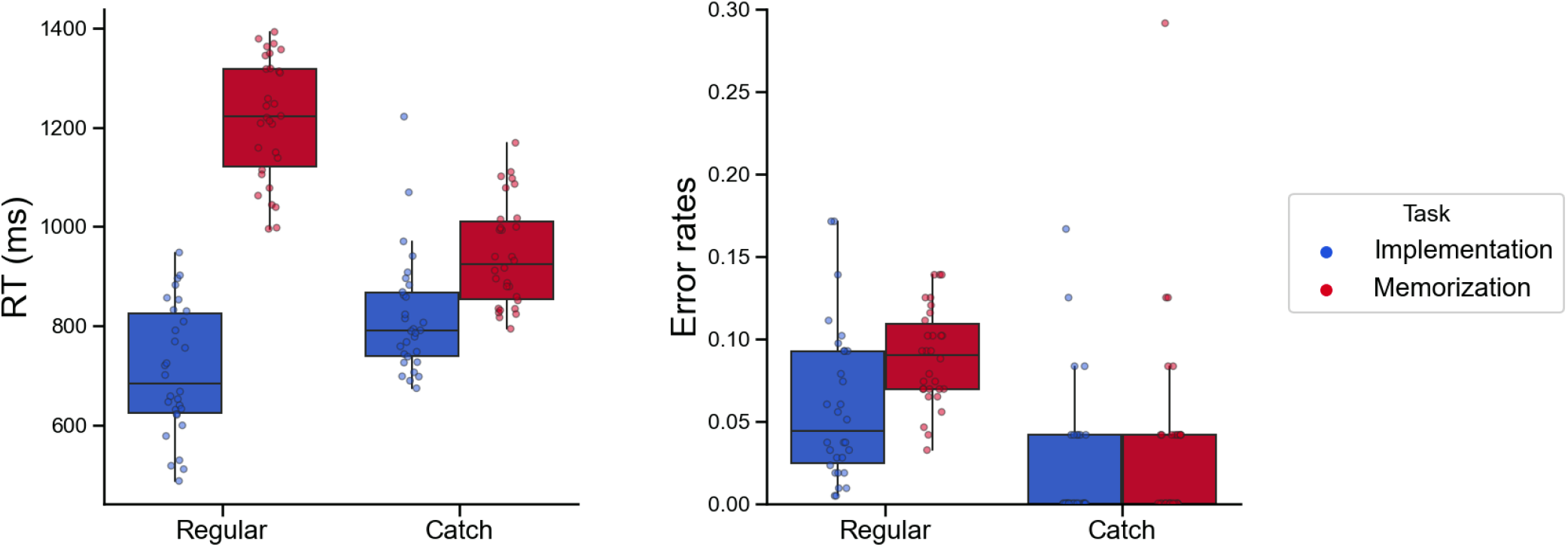
Behavioral results. Left panel: Reaction times (ms). Right panel: Error rates. In each boxplot, the thick line inside box plots depicts the second quartile (median) of the distribution (n = 30). The bounds of the boxes depict the first and third quartiles of the distribution. Whiskers denote the 1.5 interquartile range of the lower and upper quartile. Dots represent individual subjects’ scores.

### Attentional contralateral alpha suppression

To test our hypothesis that posterior alpha power tracks the orienting of attention towards the selected hemifield analogously in the two tasks, we performed a rmANOVA on the power time courses we extracted from the LatOcc ROIs, with factors Tasks (Implementation vs Memorization) and Laterality (Contralateral vs Ipsilateral to the attended hemispace). As predicted, we observed a strong reduction in alpha power contralateral to the attended hemispace (main effect of Laterality, P < 0.001, cluster corrected, d = 1.37). Importantly, no cluster of differences emerged when contrasting the two Tasks, nor when testing for the interaction of the two factors (Figure 4).

**Figure 4:**
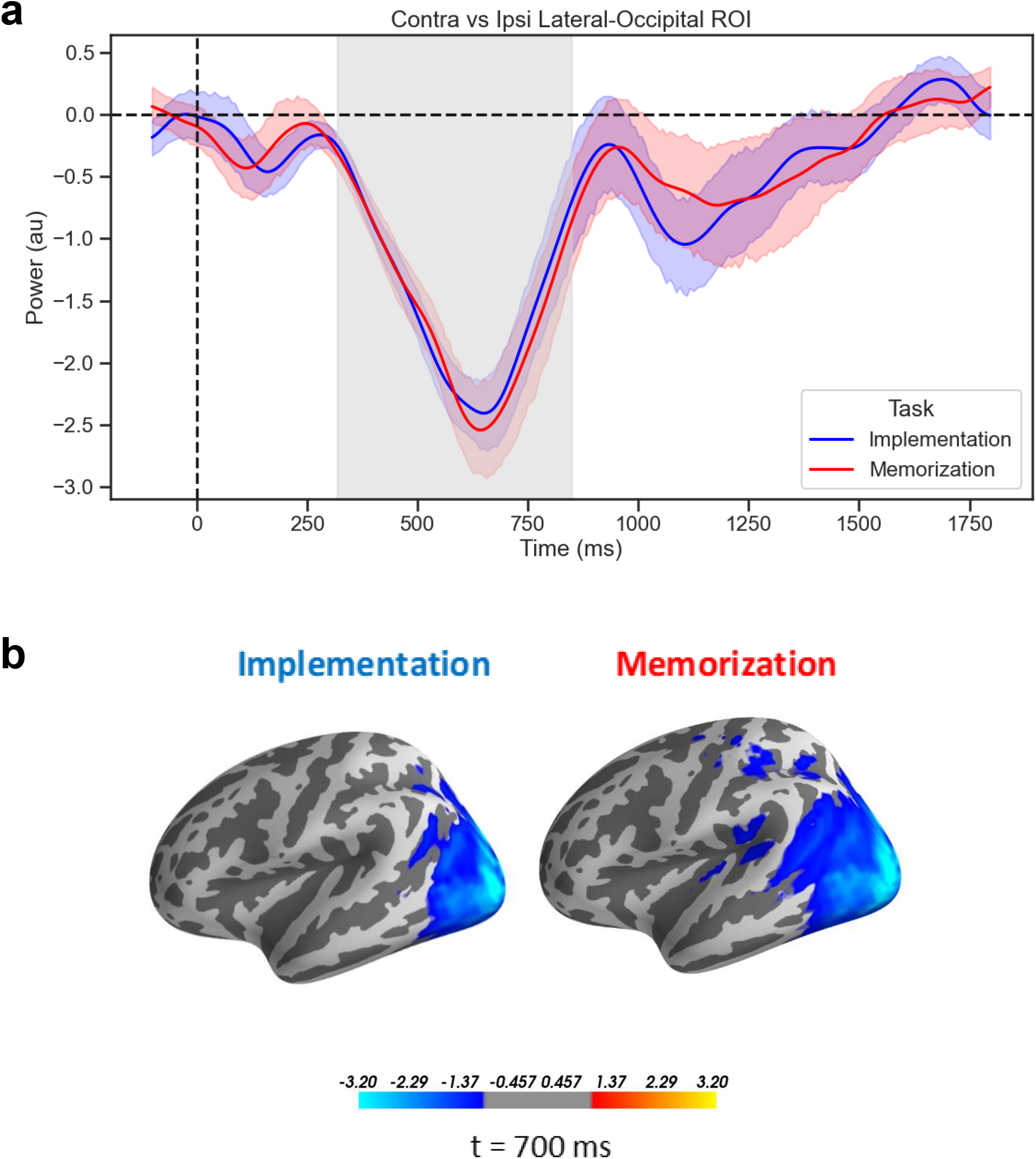
Attentional contralateral alpha suppression. **a**) Time courses of the difference waves (Contralateral vs Ipsilateral) of alpha power from the LatOcc ROIs. Shading indicates the standard error of the mean (s.e.m), gray area refers to the extent of the significant cluster for the effect of Laterality (P < 0.001, cluster corrected). **b**) Source-reconstructed activity of the alpha power for the difference between contra-and ipsilateral Cued Side, at 700 ms, for visualization purposes.

### Motor contralateral beta suppression

Activation of a specific motor plan is a crucial component of the preparation to implement novel SRs. This will then be reactivated in a reflex-like manner upon stimulus presentation (Liefooghe et al., 2012). In line with this assumption, we expected beta suppression over motor cortices to clearly track the response hand involved in the selected mappings, and particularly in the Implementation task. In other words, we predicted the time courses of beta power from the Hand ROIs to show a significant interaction in a rmANOVA with factors Tasks (Implementation vs Memorization) and Laterality (Contralateral vs Ipsilateral to the instructed response hand). We found a large significant cluster for the main effect of Laterality (P = 0.002, cluster corrected, d = 0.69). Additionally, we tested for the directional effect of the interaction with a one-sided t-test (i.e., Implementation < Memorization), and obtain a cluster with P = 0.077 (d = 0.49). No cluster was found when testing for the main effect of Task (Figure 5).

**Figure 5:**
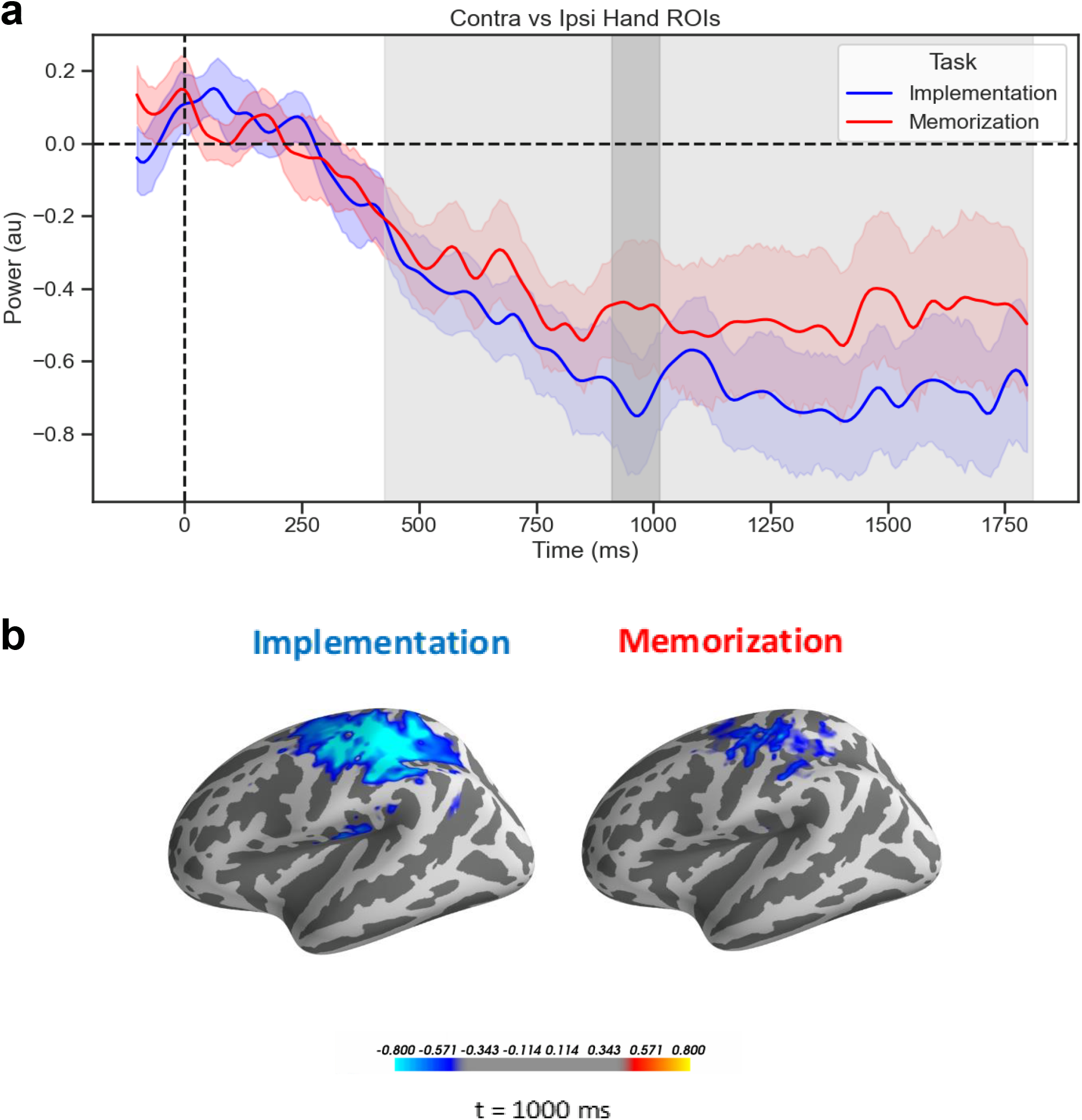
Motor contralateral beta suppression. **a**) Time courses of the difference waves (Contralateral vs Ipsilateral) of beta power from the Hand ROIs. Shading indicates the standard error of the mean (s.e.m), light gray area refers to the extent of the significant cluster for the effect of Laterality (P = 0.002, cluster corrected), dark gray area refers to the cluster for the interaction of Laterality and Task (P = 0.077, cluster corrected). **b**) Source-reconstructed activity of the beta power for the difference between contra- and ipsilateral Response Side, at 1000 ms, for visualization purposes.

### Task-specific theta increase

In our previous experiment, the amplitude of midfrontal theta oscillations significantly differed between Implementation and Memorization. To replicate this finding in the current dataset at the source level, we compared time courses of theta power from bilateral mPFC ROIs across the two tasks. Specifically, we used cluster-based permutation to test for larger theta power amplitude in Implementation compared to Memorization. We observed a significant cluster (P = 0.035, cluster corrected, d = 0.57) (Figure 6).

**Figure 6:**
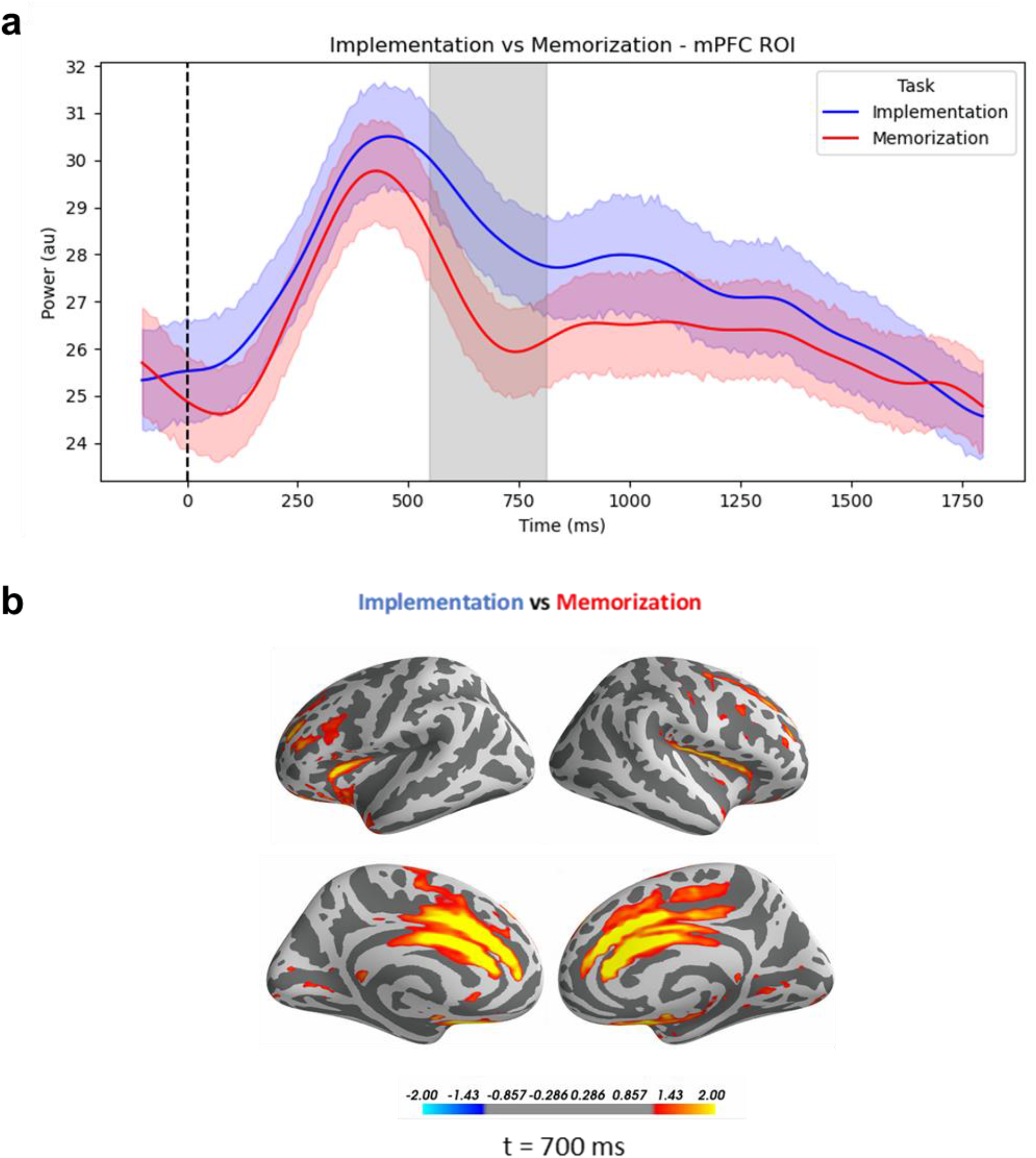
Task-specific theta increase. **a**) Time courses of theta power form the mPFC ROIs. Shading indicates the standard error of the mean (s.e.m), gray area refers to the extent of the significant cluster for the effect of Task (P = 0.035, cluster corrected). **b)** Source-reconstructed activity of theta power for the difference between Implementation and Memorization, at 700 ms, for visualization purposes.

Acitivity in the medial wall of the PFC and oscillations in the theta frequency range have been both associated with handling conflict (Cavanagh and Frank, 2014; Cavanagh and Shackman, 2015). To rule out the possibility that mPFC activity reflects the conflict elicited by incongruent conditions (i.e., trials in which the retro-cue instructed participants to orient attention towards SRs on one side of the screen and to prepare responding with the opposite hand; as compared to trialswith same Cued and Response Side), that could be larger specifically in the Implementation task, we performed additional control analyses. Namely, we compared differences in theta in congruent and incongruent trials, separately for Implementation (no clusters observed), Memorization (P = 0.168, cluster-corrected, d = 0.46), and across tasks (no clusters observed). Moreover, the effect of Congruency did not differ between the two Tasks (no clusters observed). Additonally, we tested for the effect of Task separately for congruent and incongruent trials. While there was a significant difference between Tasks in congruent trials (P = 0.033, cluster-corrected, d = 0.54), this was not the case for incongruent trials (P = 0.124, cluster-corrected, d = 0.43). Although these results cannot be taken as evidence for an interaction between the factors Task and Congruency (Nieuwenhuis et al., 2011), they are hinting at a larger difference in theta oscillations between Tasks in congruent trials (i.e., trials eliciting less conflict). Taken together, these exploratory analyses support our interpretation that the observed mPFC activity is indicative of differences between Tasks rather than the by-product of conflict resolution.

### mPFC theta power mediates the effect of Task on RTs

We hypothesized that trial-by-trial theta power in the mPFC mediated the effect of Task on behavioral performance. First, we tested for an effect of Task on RTs, by fitting a LMM with a fixed effect of Task, a random slope for Task, and a random intercept for each participant (*RT ∼ Task + (1 + Task | Subject)).* As expected, Implementation was associated with significantly faster RTs (t_28.83_ = -28.75, β = -257.50, CI 95% = [-275.05, - 239.94], p < 0.001). Moreover, by fitting a LMM with identical structure to predict theta power (*Mean_Theta ∼ Task + (1 + Task | Subject)),* we showed that single-trial variations in mPFC theta power were significantly dependent upon task demands (t_28.83_ = 2.72, β = 0.10, CI 95% = [0.03, 0.18], p = 0.011), such that larger theta values were associated with the Implementation task.

Finally, we also tested for the fixed effects of both Task and theta power (while assuming random slope for Task and random intercept for each participant) by fitting the LMM *RT ∼ Mean_theta + Task + (1 + Task | Subject)*. We found a significant effect of theta power t_11200.23_ = -3.54, β = -3.29, CI 95% = [-5.11, -1.47], p < 0.001, suggesting faster RTs for larger theta values. The effect of Task was significant, t_28.83_= -28.70, β = -257.16, CI 95% = [-274.72, -239.60], p < 0.001. Note that adding an interaction term as fixed effect to this model (i.e., Mean_Theta * Task) did not result in a significant interaction effect (p = 0.66). (Figure 7a).

**Figure 7:**
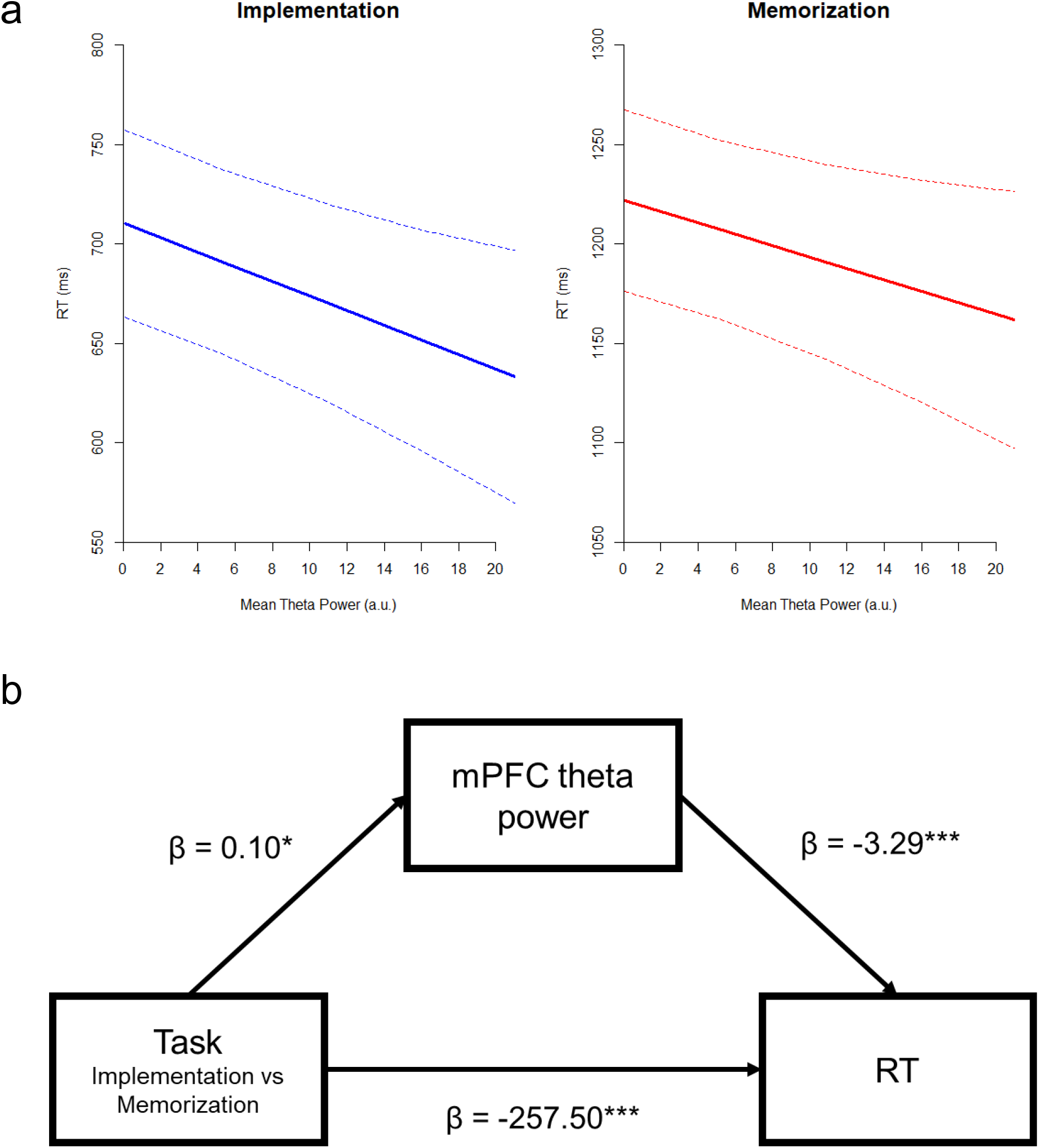
Effect of mean theta power on RTs. Panel a): Effect of theta power on RTs, for both Implementation and Memorization. High values of theta power are associated with faster RTs in both tasks. The interaction between theta power and Task was not significant. The dotted lines show the 95% confidence intervals. Panel b): Mediation model with beta values. Task significantly influenced mPFC theta power, which in turns affected RTs. Therefore, theta power mediates the effect of tasks demands on behavioral performance. However, the direct effect of Task on RTs remained significant also when accounting for the mediating influence of theta power, suggesting a partial mediation.

This pattern of results suggests that theta partially mediated the effect of task, and this was directly tested by a causal mediation analysis revealing not only a significant direct effect of Task on RTs, β = 514, CI 95% = [477, 549.73], p < 0.001, but also a significant indirect effect via theta power, β = 0.66, CI 95% = [0.11, 1.42], p = 0.013, indicative of a partial mediation (Figure 7b).

### Functional connectivity between mPFC and motor/visual areas

To test our hypothesis that preparing to implement SRs is characterized by increased connectivity in theta band between frontal and motor areas, we used LMM to estimate whether different task demands resulted in different PLV values between mPFC and Hand ROIs. We fitted a model with a fixed effect for Task, Laterality and their interaction, a random effect for Task and a random intercept for each participant (*PLV(mPFC-Hand) ∼ Task * Laterality + (1 + Task | Subject))*. We found a significant effect of Task (t0.29 = 3.04, β = 0.008, CI 95% = [0.0003, 0.001], p = 0.005), thus proving stronger connectivity during Implementation compared to Memorization (Figure 8). Contrary to our hypothesis, the effect of Laterality and its interaction with Task were not significant (p = 0.16 and p = 0.71, respectively).

**Figure8:**
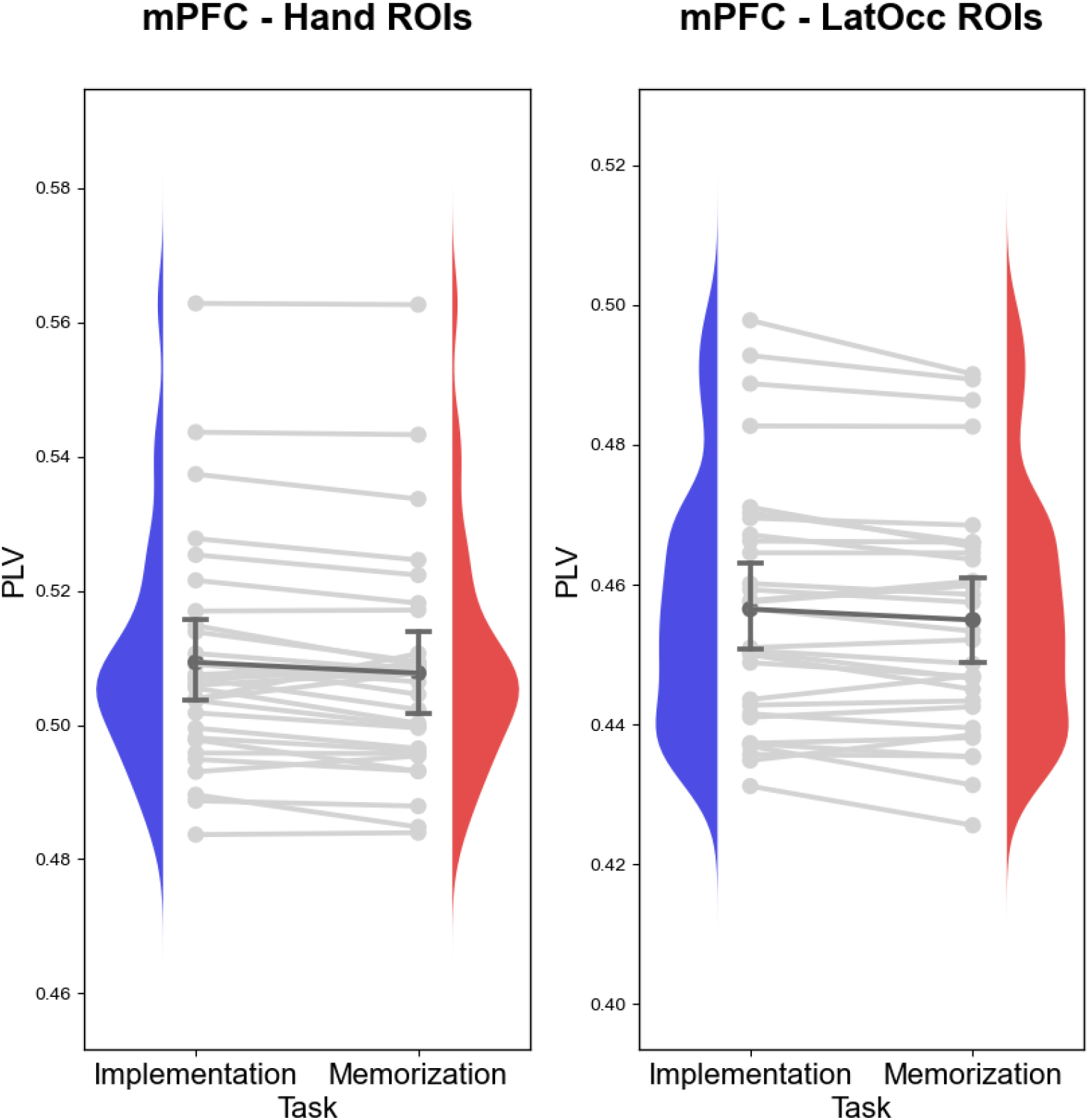
Connectivity between mPFC and posterior ROIs. Linear mixed effect models revealed that Implementation task-demands are associated with stronger connectivity in the theta frequency range between mPFC and motor regions (left panel) and visual regions (right panel). For visualization purposes, the plots depict subject-level averages. Blue and red curves represent the density distributions of subject-level averages of PLV of Implementation and Memorization, respectively. Light gray lines connect the average in the two Tasks for each individual participant, whereas the dark gray line connect the group-level averages (whiskers denote 95% confidence intervals).

Analogously, we tested whether task demands affected the degree of connectivity between frontal and visual areas. We fitted a model with the structure *PLV(mPFC-LatOcc) ∼ Task * Laterality + (1 + Task | Subject*). We found a significant effect of Task (t0.29 = 2.96, β = 0.008, CI 95% = [0.0002, 0.001, p = 0.006), supporting our hypothesis of stronger connectivity during Implementation compared to Memorization (Figure 8). Again, the effect of Laterality and the interaction of Laterality and Task were not significant (p = 0.38 and p = 0.85, respectively).

## Discussion

The present study aimed at providing insights in the inter-areal dynamics supporting instructions following, and to replicate our previous results highlighting implementation-specific neural oscillations. This line of work is motivated by the assumption that the intention to implement the instructed SRs triggers a cascade of cognitive processes, ultimately leading to the emergence of a representational state intrinsically different from the initially encoded symbolic content (Muhle-Karbe et al., 2017; González-García et al., 2021).

Such reformatting of prioritized items into a *behavior-optimized format* has been recently proposed as core mechanism to efficiently deal with multiple memoranda and circumvent capacity limitations(Myers et al., 2017.

In the context of instructions implementation, we hypothesized that this optimized format consists of strengthened connections between sensory and motor areas representing task-relevant information, coordinated by medial prefrontal structures through long-range phase synchronization. This operationalization is grounded in a body of computational work addressing the role of mPFC theta oscillations in flexibly binding task-relevant areas for upcoming task-demands (Verguts, 2017; Verbeke and Verguts, 2019; Senoussi et al., 2020b; Verbeke et al., 2020).

In line with our previous results (Formica et al., 2021), in both tasks we found a sharp suppression of alpha oscillations in the lateral occipital area contralateral to the attended hemispace. This feature, analogous across task-demands, is a hallmark of attentional resources deployment, with the putative function of gating information inflow and contributing to the creation of a functional network (Sauseng et al., 2005; Jensen and Mazaheri, 2010; Mazaheri et al., 2014; Poch et al., 2017; van Ede, 2017; Van Diepen et al., 2019; Keefe and Störmer, 2021). This finding suggests that the two tasks do not differ with respect to resource allocation.

On the contrary, neural signatures of motor preparation were expected to be more prominent during Implementation, given the possibility to activate a specific action plan. We focused on beta dynamics in the motor cortex (Pfurtscheller and Neuper, 1997; Cheyne, 2013; Schneider et al., 2017) and found in both tasks a sustained suppression of beta oscillations contralateral to the instructed response hand, indicating the lateralization of motor preparation. Notably, and contrary to our hypothesis, this suppression was not task-specific, and only marginally larger in Implementation. Although unexpected, this finding might be highlighting the recruitment of motor areas also for the declarative maintenance of SRs that will never have to be overtly executedsupporting the idea of distributed and content-specific maintenance substrates in WM (Christophel et al., 2017).

A second crucial oscillatory feature associated with the proactive preparation for SRs implementation is an increase in midfrontal theta power (Formica et al., 2021). This has been identified as a spectral signature shared across a plethora of mechanisms involved in adaptive control (Cavanagh and Frank, 2014; Cavanagh and Shackman, 2015), with neural generators in the mPFC (De La Vega et al., 2016). Accordingly, in the current study we observed significantly larger mPFC theta oscillations during Implementation compared to Memorization. Importantly, we highlighted the specific functional relevance of this neural activity by showing how theta power mediates the effect of different task-demands on behavioral performance (Bridwell et al., 2018).

Stemming from this literature, mPFC has been attributed a central role in determining and instantiating control policies in several computational models of proactive and reactive cognitive control (Dosenbach et al., 2008; Botvinick and Cohen, 2014; Shenhav et al., 2017; Verguts, 2017; Holroyd and Verguts, 2021). In particular, in the work of Verguts and colleagues, mPFC theta oscillations signal the need for adjustments to reach the current goal (Verguts, 2017; Verbeke and Verguts, 2019; Senoussi et al., 2020b; Verbeke et al., 2020), and achieve them by synchronizing the activity of task-relevant pairs of sensory and action units, thereby allowing them to communicate more efficiently (Fries, 2005, 2015). Therefore, these models identify in the mPFC the structure responsible to coordinate and operationalize the flexible binding of lower-level modules to meet task-demands.

Our theta phase connectivity hypotheses are embedded within this framework, insofar implementation task-demands are thought to require the coordination of sensory and motor information in a coherent action-oriented representation. Accordingly, we found that theta oscillations between mPFC and motor/visual areas were significantly more synchronized in the Implementation task. Crucially, these findings show that proceduralization triggers the formation of a functional network encompassing frontal and posterior areas through synchronization, compatibly with a view of mPFC exerting top-down control towards posterior areas in response to the specific task-demands.

Notably, contrary to our hypotheses, motor/visual areas contralateral and ipsilateral to the hand and hemispace currently relevant showed no difference in their degree of connectivity to the mPFC. This finding is open to alternative explanations. First, it might suggest that top-down synchronization is exerted similarly towards both hemispheres independently of lateralization of stimuli and responses. This interpretation is consistent with a previous study testing the model prediction in a reversal rule learning task involving lateralized responses and reporting bilateral clusters of connectivity between FCz and posterior electrodes (Verbeke et al., 2020). Information on which networks are task-relevant and which should be inhibited could be coded in other aspects of the interactive dynamics between mPFC and posterior areas, such as phase coding or cross-frequency coupling (Helfrich and Knight, 2016), or by means of activity-silent and less resource consuming neurophysiological mechanisms (Stokes, 2015; Masse et al., 2019).

Alternatively, the lack of a significant lateralization effect might be attributed to the selected spatial and temporal features. It cannot be ruled out, for instance, that a short-lasting lateralization in the connectivity patterns emerges only late during the cue-target interval. However, this explanation appears unlikely given the early-onset suppression of beta oscillations in the motor cortex contralateral to the selected response hand, indicating that information on the specific effector is already present shortly after the retro-cue. Moreover, although the selected ROIs were suited to detect the hypothesized lateralized local changes in oscillatory activity, it is possible that lateralization in connectivity with the mPFC is implemented at different stages of the processing hierarchy. Connectivity between mPFC and premotor, rather than motor, cortices might carry information on the currently relevant response hand, and, in a similar fashion, areas at later stages of the visual stream might be more synchronized depending on the stimulus material. In this regard, recent fMRI evidence showed connectivity between the anterior cingulate cortex and visual areas specific for the processing of faces and houses, scaling with task difficulty (Aben et al., 2020). This finding hints at the need for a more fine-grained ROIs definition, and further research should investigate this issue for instance by using independent functional localizers (Baldauf and Desimone, 2014; Kok et al., 2017; Senoussi et al., 2020a; González-García et al., 2021).

In the model of flexible binding, the end point is the synchronization of the posterior brain areas coding for the currently relevant stimulus and response. This would be implemented in the gamma frequency range, with the bursts aligning the phase of gamma oscillations (Verguts, 2017). Such synchronization between processing units is difficult to test in the present dataset, as high frequencies (> 30 Hz) are significantly more difficult to investigate with EEG (Nottage and Horder, 2015). Although not directly providing evidence that procedural representations consist of the binding of stimulus and response information, our results support the hypothesis that implementation task-demands elicit the emergence of a strengthened network of task-relevant brain areas coordinated by the mPFC and instantiated by means of theta-phase synchronization.

Importantly, while theta oscillations from the mPFC provide an explanation on *how* posterior areas achieve synchronization, the information on *which* regions should become synchronized is thought to be coded in the lateral PFC (Verguts, 2017). This is consistent with fMRI literature, showing lateral PFC to be a pivotal region in task-set coding (Duncan, 2001; Woolgar et al., 2011; Shahnazian et al., 2021) and dissociating between maintenance and execution task demands (Demanet et al., 2016; González-García et al., 2017, 2021; Muhle-Karbe et al., 2017; Palenciano et al., 2019a). Here, we put forward and tested hypotheses on theta activity in the mPFC, as these were clearly derivable from the literature, while remaining agnostic to the role of the lateral PFC. Further research should clarify the oscillatory phenomena associated with instructions following in the lateral PFC, contributing to integrate the results from the two techniques.

In summary, we focused on the role of mPFC theta oscillations as core mechanism to coordinate the communication between brain areas relevant to execute the instructed behavior. We showed that proactively preparing to implement novel SRs elicits an increase in theta-phase synchronization between frontal control areas and motor/visual areas, supporting the idea that a procedural, action-oriented format relies on the interplay of distant brain regions.

The authors declare no competing financial interests.

## Acknowledgements

The present study was funded by Special Research Fund of Ghent University BOF.GOA.2017.0002.03 and the Deutsche Forschungsgemeinschaft (DFG, German Research Foundation) under Germany’s Excellence Strategy – EXC 2002/1 “Science of Intelligence” – project number 390523135.

C.G.G. was additionally funded by European Union’s Horizon 2020 research and innovation programme under the Marie Sklodowska-Curie grant agreement no. 835767, and the Spanish Ministry of Science and Innovation, grant ID IJC2019-040208-I.

M.S. was supported by Research Foundation Flanders (FWO) grant number G012816N, and the Research Council of Ghent University, grant BOF17-GOA-004.

M.B. is supported by an Einstein Strategic Professorship (Einstein Foundation Berlin).

The authors wish to thank Dr. Ricardo Bruña and Dr. Emiel Cracco for the fruitful discussions, insightful comments, and kind support during data analysis.

## Authors contribution

All authors designed the study. S.F. collected and analyzed the data, which were then interpreted together with all authors. S.F. wrote the manuscript and all authors were involved in the revisions. M.B supervised this work.

## Supplementary Materials

### 1. Exploratory three-way ANOVAs on RTs and Error Rates

The exploratory three-way ANOVA on RTs confirmed the main effect of Task (*F*_29,1_ = 841.59, p < 0.001, η²p = 0.98) and additionally yielded significant effects of Response Side (*F*_29,1_ = 4.74, p = 0.038, η²p = 0.14) and the interaction of Cued * Response Side (*F*_29,1_ = 6.19, p = 0.019, η²p = 0.18). The corresponding ANOVA on Error Rates showed significant effects for Task (*F*_29,1_ = 13.36, p < 0.001, η²p = 0.31), Response Side (*F*_29,1_ = 5.64, p = 0.024, η²p = 0.16), the interaction of Cued * Response Side (*F*_29,1_ = 6.50, p = 0.016, η²p = 0.18), and the three-way interaction of Task * Cued Side * Response Side (*F*_29,1_ = 7.66, p = 0.010, η²p = 0.21).

**Figure S1:**
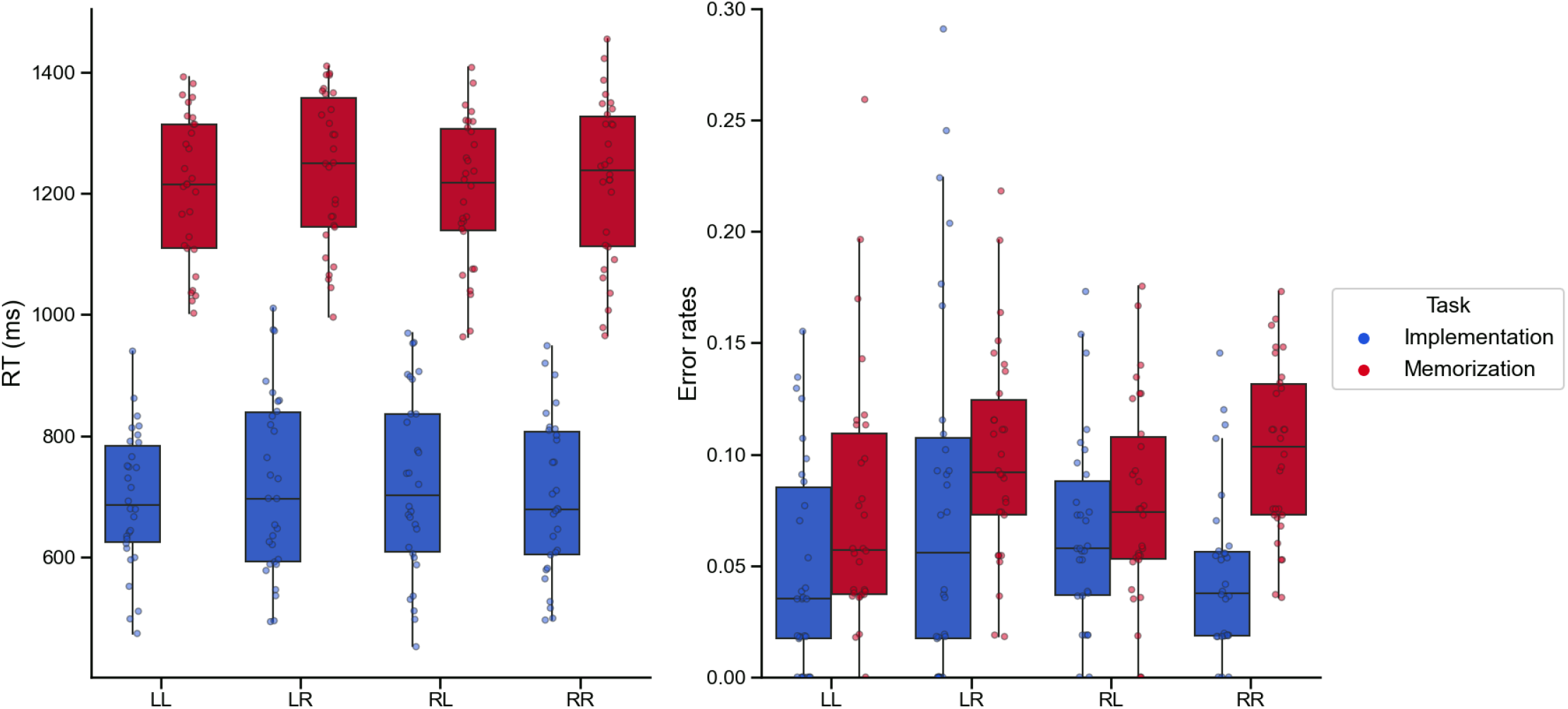
Exploratory three-way ANOVAs on RTs and Error rates. Left panel: Reaction times (ms). Right panel: Error rates. X-axis labels refer to the individual conditions resulting from the crossing of Cued Side and Response Side. The first letter indicates the cued hemispace (L for Left and R for right), the second letter denotes the instructed response hand (L for Left and R for Right). In each boxplot, the thick line inside box plots depicts the second quartile (median) of the distribution (n = 30). The bounds of the boxes depict the first and third quartiles of the distribution. Whiskers denote the 1.5 interquartile range of the lower and upper quartile. Dots represent individual subjects’ scores.

## Footnotes

1 Importantly, similar results were obtained without applying any filtering procedure to the data.

